# Targeted protein degradation via CAR endocytosis of antigen in T cells

**DOI:** 10.1101/2024.11.20.624443

**Authors:** Youguang Wang, Na Yin, Min Peng

## Abstract

Targeted protein degradation (TPD) relies on molecules engaging host protein degradation machinery. Here, we developed a novel TPD platform based on antigen endocytosis and degradation by chimeric antigen receptor (CAR) T cells. T cells expressing a CAR with TNFR1 as the antigen-binding domain (TNFR1T) were able to bind, endocytose, and degrade TNF in vitro. To enhance in vivo expansion and persistence of TNFR1T cells, BCOR and ZC3H12A were depleted, generating TNFR1T_IF_ cells. In a human TNF transgenic mouse model of rheumatoid arthritis, a single infusion of TNFR1T_IF_ cells sustainably reduced serum hTNF to near wild-type levels, leading to long-term disease remission. This approach extends CAR T cell targets from cells to extracellular proteins, enabling long-term degradation of inflammatory cytokines and durable remission in chronic inflammatory diseases.

## Introduction

Extracellular factors, such as cytokines, are implicated in various diseases. Utilizing biologics to functionally neutralize these extracellular factors is a valid therapeutic approach in the clinic. While biologics have shown effectiveness in disease treatment, they have inherent limitations that require further improvement. Due to their short half-lives, biologics must be administered repeatedly to maintain therapeutic efficacy, leading to increased costs, reduced patient compliance, and decreased quality of life. For example, the widely used anti-TNF antibody (Adalimumab, Humira) requires bi-monthly injections for indications like rheumatoid arthritis (RA)(Colombel et al., 2007; Menter et al., 2008; Weinblatt et al., 2003). Repeated protein injections also cause side effects, including injection site reactions, anaphylaxis, etc.(Kim et al., 2023; Weinblatt et al., 2003), and the development of anti-drug antibodies that eventually diminish therapeutic efficacy(de Speville and Moreno, 2021; Ridker et al., 2017; Wu et al., 2016).

Targeted protein degradation (TPD) technologies, such as proteolysis-targeting chimeras (PROTACs), offer advantages over functional inhibition via biologics(Dale et al., 2021). While traditional PROTACs can only target intracellular proteins for degradation, newer TPD approaches, like LYTACs(Banik et al., 2020), AbTACs(Cotton et al., 2021), ProTABs(Marei et al., 2022), and KineTACs(Pance et al., 2023), among others(Caianiello et al., 2021; Hamada et al., 2023; Sun et al., 2023; Zhang et al., 2021), can target membrane and/or extracellular proteins by engaging surface E3 ubiquitin ligases and/or the endocytosis-lysosome pathway. However, few of these novel approaches have demonstrated potent and durable therapeutic effects in disease models in vivo. Furthermore, existing TPD methods only induce transient protein degradation, making them unsuitable for chronic diseases that require long-term treatment.

All existing TPD technologies rely on engaging the endogenous protein degradation machinery of host cells(Békés et al., 2022; Schapira et al., 2019). For instance, insulin growth factor 2 receptor (IGF2R, also known as CI-M6PR) has been widely used as an endocytosis receptor to trigger targeted degradation of membrane or extracellular proteins in vitro(Banik et al., 2020; Mikitiuk et al., 2023; Zhang et al., 2023). However, IGF2R mutant mice exhibit excessive IGF2 signaling via IGF1R, resulting in perinatal lethality(Ludwig et al., 1996). This indicates that the physiological degradation of IGF2 by IGF2R is critical for maintaining an appropriate level of IGF2 in vivo. Forced endocytosis of IGF2R by exogenous agents in vivo could interfere with the physiological degradation of IGF2, potentially causing side effects. Similarly, other commonly used endocytosis receptors and surface E3 ubiquitin ligases, such as transferrin receptor (TfR) and RNF43/ZNFR3(Marei et al., 2022; Su et al., 2024), also actively endocytose and degrade their native ligands(de Lau et al., 2014; Hao et al., 2012; Huebers and Finch, 1987; Koo et al., 2012). Redirecting these receptors for TPD likely interferes with the degradation of their native substrates. Indeed, unforeseeable side effects have been observed in clinical trials of TPD drugs(Duyvendak et al., 2005; Hamadani et al., 2007; Merrill et al., 2022; Richardson et al., 2023; Sohlbach et al., 2006). Thus, engaging host protein degradation machinery is an inherent flaw present in all current TPD technologies. Additionally, the effects of current TPD technologies heavily depend on the expression pattern and level of E3 ubiquitin ligases or endocytosis receptors, limiting their in vivo efficacy. All these limitations highlight the need for developments that bypass the engagement of endogenous protein degradation machinery.

CAR T cells, as an exogenous entity, traditionally target cells rather than soluble factors(Finck et al., 2022; Irvine et al., 2022; Labanieh and Mackall, 2023; Sadelain et al., 2017). In this study, we engineered CAR T cells as an exogenous platform to bypass host protein degradation machinery for the targeted and long-term degradation of an extracellular factor: the inflammatory cytokine TNF.

## Results

### TNFR1 CAR T (TNFR1T) cells endocytose TNF in vitro

To target TNF, we designed a second-generation CAR with the extracellular domain (amino acids 1-212) of mouse TNF receptor 1 (TNFR1) as the antigen-binding domain, followed by the CD28 transmembrane/intracellular domain and CD3ζ chain (Figure 1A). Given that mouse TNFR1 binds to both mouse and human TNF(Lewis et al., 1991), we could evaluate the responses of TNFR1 CAR T (TNFR1T) cells to both mouse and human TNF using the same CAR (Figure 1B). TNFR1 CAR expression was coupled with a Thy1.1 marker for the detection and purification of CAR T cells (Figure 1C). Co-staining T cells transduced with a retrovirus expressing this TNFR1 CAR with anti-Thy1.1 and anti-TNFR1 antibodies revealed surface expression of TNFR1 CAR on T cells (Figure 1D), with HER2 CAR T cells serving as controls. In an in vitro binding assay, an RFP-TNF fusion protein demonstrated specific binding to TNFR1T cells but not HER2 CAR T cells (Figure 1E). These data demonstrate that TNFR1 CAR is expressed on T cells and binds to soluble TNF.

**Figure 1.**
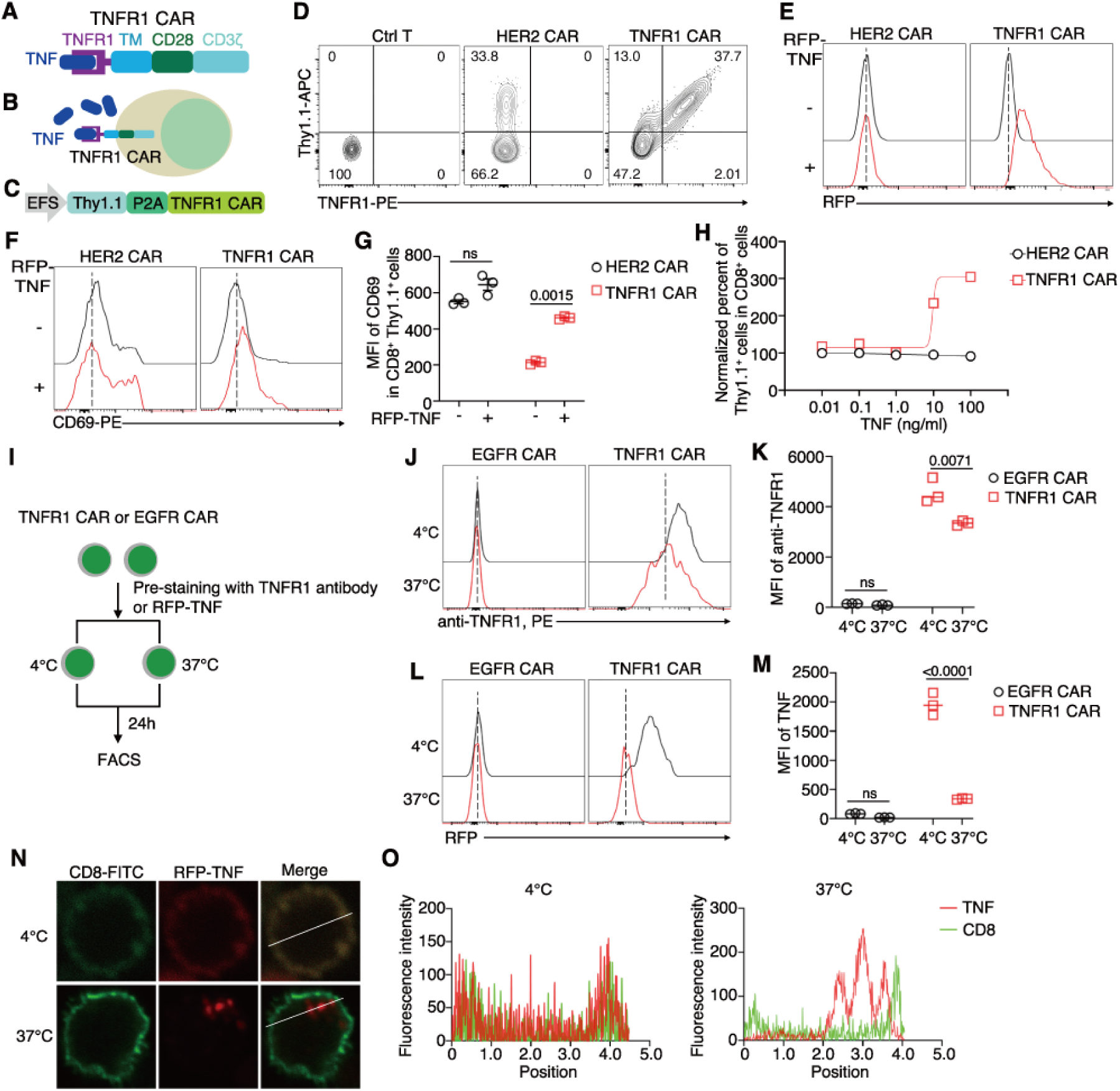
TNFR1T cells endocytose TNF in vitro. **A**, Design of TNFR1 CAR. **B**, Illustration depicting the interaction between TNF and TNFR1T cells. **C**, Vector for co-expression of TNFR1 CAR and a Thy1.1 marker. **D**, Expression of Thy1.1 and the indicated CAR on T cells. Representative plots from two independent experiments are shown. **E**, Binding of RFP-TNF to the indicated CAR T cells. Representative plots from two independent experiments are shown. **F**, **G**, Expression of CD69 on the indicated CAR T cells after 24-hour stimulation with RFP-TNF. Representative plots (**F**) and statistics (**G**) are shown (n = 3 independent experiments). **H**, Relative expansion of CAR T cells in the presence of indicated concentrations of recombinant TNF. Representative data from two independent experiments are shown. **I**, Experimental design for TNF internalization assay. **J** - **M**, Flow cytometry analysis of anti-TNFR1 signals (**J**, **K**) and RFP-TNF signals (**L**, **M**) in TNFR1T cells. Representative plots (**J**, **L**) and statistics (**K**, **M**) are shown (n=3 independent experiments). **N**, Immunofluorescent examination of RFP-TNF localization in TNFR1T cells kept at 4℃ or after 24-hour incubation at 37℃. Representative images are shown. **O**, Quantification of fluorescent signals along lines drawn in cells shown (**N**). **G**, **K**, **M**, Data represent mean ± SEM from one of three independent experiments. *P* values are shown, ns, not significant, two-tailed unpaired Student’s t-test.

Upon incubation with RFP-TNF, TNFR1T cells, but not HER2 CAR T cells, exhibited an upregulation of CD69 (Figure 1F and 1G), an activation marker of T cells, indicating that soluble TNF activates TNFR1T cells. Consistently, recombinant native mouse TNF (not the RFP-TNF fusion protein) enhanced the expansion of TNFR1T cells, but not HER2 CAR T cells in vitro (Figure 1H). These data demonstrate that soluble TNF specifically activates and promotes the expansion of TNFR1T cells in vitro.

CAR is known to undergo endocytosis in T cells(Li et al., 2020). To test whether CAR endocytosis can be leveraged to internalize its antigen, we utilized an in vitro endocytosis assay (Figure 1I). In this assay, TNFR1T cells or control HER2 CAR T cells were first labeled with anti-TNFR1 antibody or RFP-TNF at 4℃. After washing out unbound antibody or RFP-TNF, CAR T cells were resuspended in T cell medium and split into two groups: one group kept at 4℃ and the other incubated at 37°C for endocytosis (Figure 1I). After 24 hours, the signals of anti-TNFR1 antibody and RFP-TNF were reduced in TNFR1T cells kept at 37°C compared with those kept at 4℃ (Figure 1J-1M), suggesting that anti-TNFR1 antibody and RFP-TNF were either endocytosed and degraded in TNFR1T cells or dissociated from TNFR1T cells. Immunofluorescent staining showed that RFP-TNF could be detected within TNFR1T cells after incubation at 37°C (Figure 1N and 1O), indicating RFP-TNF is endocytosed by TNFR1T cells.

Together, these data demonstrate that TNFR1T cells respond to TNF and endocytose TNF in vitro.

### TNFR1T cells lack therapeutic efficacy for RA in hTNF-tg mice due to poor in vivo expansion

Since TNFR1T cells are capable of endocytosing TNF in vitro, we investigated their potential therapeutic efficacy in rheumatoid arthritis (RA), a disease that responds to anti-TNF antibodies(Weinblatt et al., 2003; Wu et al., 2016). We administered TNFR1T cells to human TNF transgenic (hTNF-tg) mice (Figure 2A), a widely used RA model(Keffer et al., 1991; Li and Schwarz, 2003). In comparison to Humira (adalimumab), an FDA-approved treatment for RA(Weinblatt et al., 2003), TNFR1T cells did not reduce the clinical severity of RA in hTNF-tg mice (Figure 2B) and failed to improve grip strength (Figure 2C). Consistent with the lack of therapeutic effect, TNFR1T cells were undetectable in hTNF-tg mice (Figure 2D). These results suggest that the failure of TNFR1T cells to expand and/or persist in vivo likely accounts for their inability to treat RA in this model.

**Figure 2.**
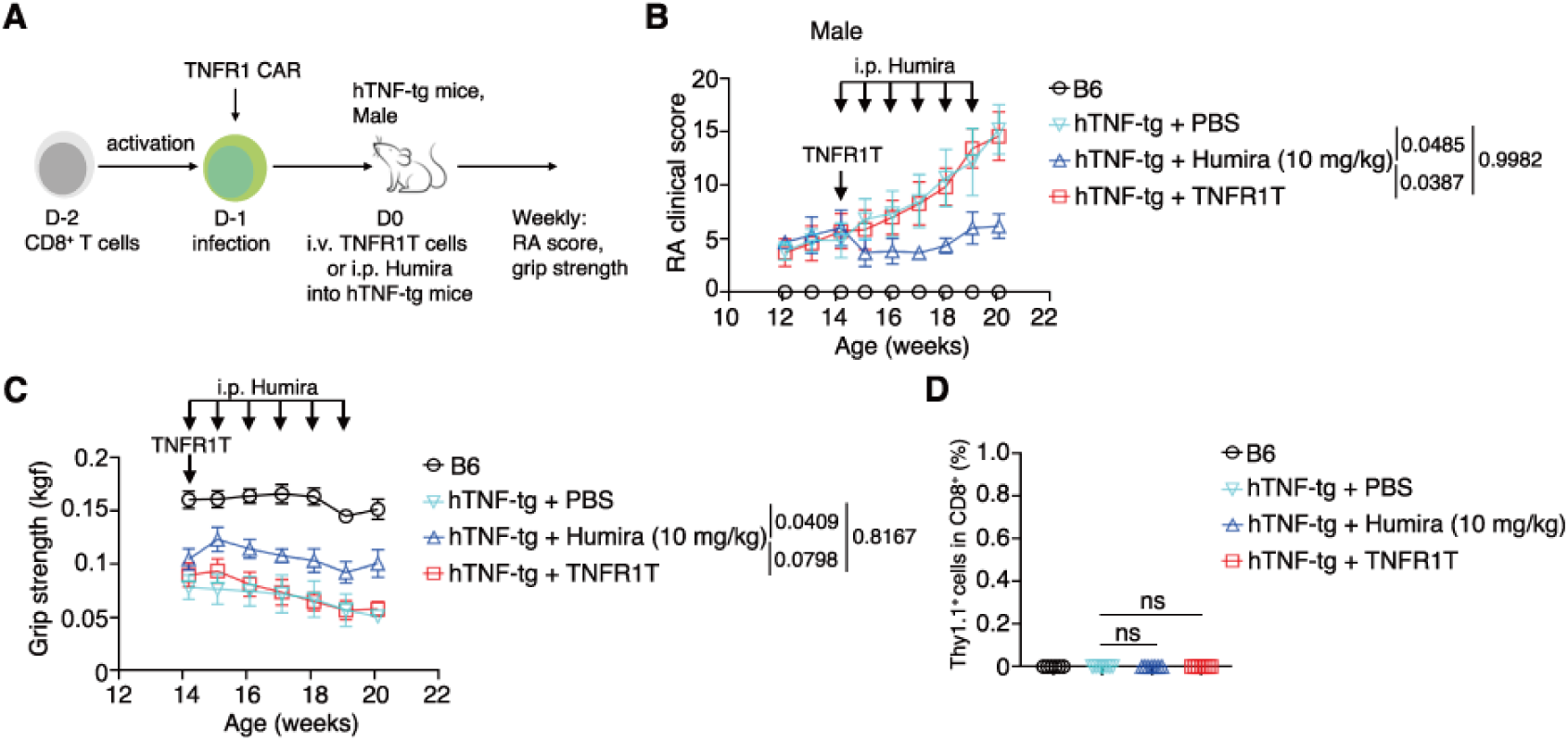
TNFR1T cells lack therapeutic efficacy for RA in hTNF-tg mice due to insufficient in vivo expansion. **A**, Experimental design for treating hTNF-induced RA with PBS, TNFR1T cells or Humira. **B**, RA clinical scores in hTNF-tg and B6 mice across treatment groups (n = 6 mice per group). **C**, Grip strength in hTNF-tg and B6 mice across treatment groups (n = 6 mice per group). **D**, Percentage of Thy1.1^+^ CAR T cells in the spleen 42 days post-transfer. **B**, **C**, **D**, Data represent mean ± SEM from one of two independent experiments. *P* values are shown, ns, not significant, two-way ANOVA multiple-comparisons test in (**B, C**), one-way ANOVA multiple-comparisons test in (**D**).

### TNFR1T_IF_ cells expand and persist in vivo

To further explore the expansion of TNFR1T cells in vivo, we transferred these cells into B6 mice (Figure S1A). No TNFR1T cells could be detected in the peripheral blood (or any organs) of recipient mice (Figure S1B and S1C, and data not shown), indicating a lack of expansion of TNFR1T cells in vivo in the immunocompetent B6 mice. These observations are consistent with a critical role of lymphodepleting pre-conditioning in the expansion of CAR-T cells in immunocompetent hosts(Dudley et al., 2002; Murad et al., 2021).

To address TNFR1T cells’ inability to expand in vivo, we employed a recently reported strategy involving the simultaneous knockout of BCOR and ZC3H12A to enhance CAR T cell expansion and persistence in immunocompetent mice without conditioning(Jin et al., 2024; Wang et al., 2024). BCOR- and/or ZC3H12A-deficient TNFR1T cells were generated by co-delivering TNFR1 CAR and sgRNAs into activated Cas9^+^CD8^+^ T cells (Figure S1D). Successful editing of *Bcor* and *Zc3h12a* genes was confirmed by DNA sequencing (Figure S1E and S1F). TNFR1T cells expressing sgRNAs targeting non-targeting (NT) control (sgNT), *Bcor* (sg*Bcor*), or *Zc3h12a* (sg*Zc3h12a*) showed no expansion in vivo (Figure 3A and 3B). In contrast, TNFR1T cells expressing sgRNAs targeting both *Bcor* and *Zc3h12a* (sg*Bcor*/*Zc3h12a*) exhibited robust expansion and persistence (Figure 3A and 3B), reflecting a synergistic effect between BCOR and ZC3H12A deficiencies in TNFR1T cell expansion and persistence in vivo, consistent with previous findings in other BCOR- and ZC3H12A-deficient CAR T cells(Jin et al., 2024; Wang et al., 2024). For simplicity, we referred to TNFR1T cells devoid of both BCOR and ZC3H12A as immortal-like and functional TNFR1T (TNFR1T_IF_) cells, following previous naming pattern(Jin et al., 2024; Wang et al., 2024).

**Figure 3.**
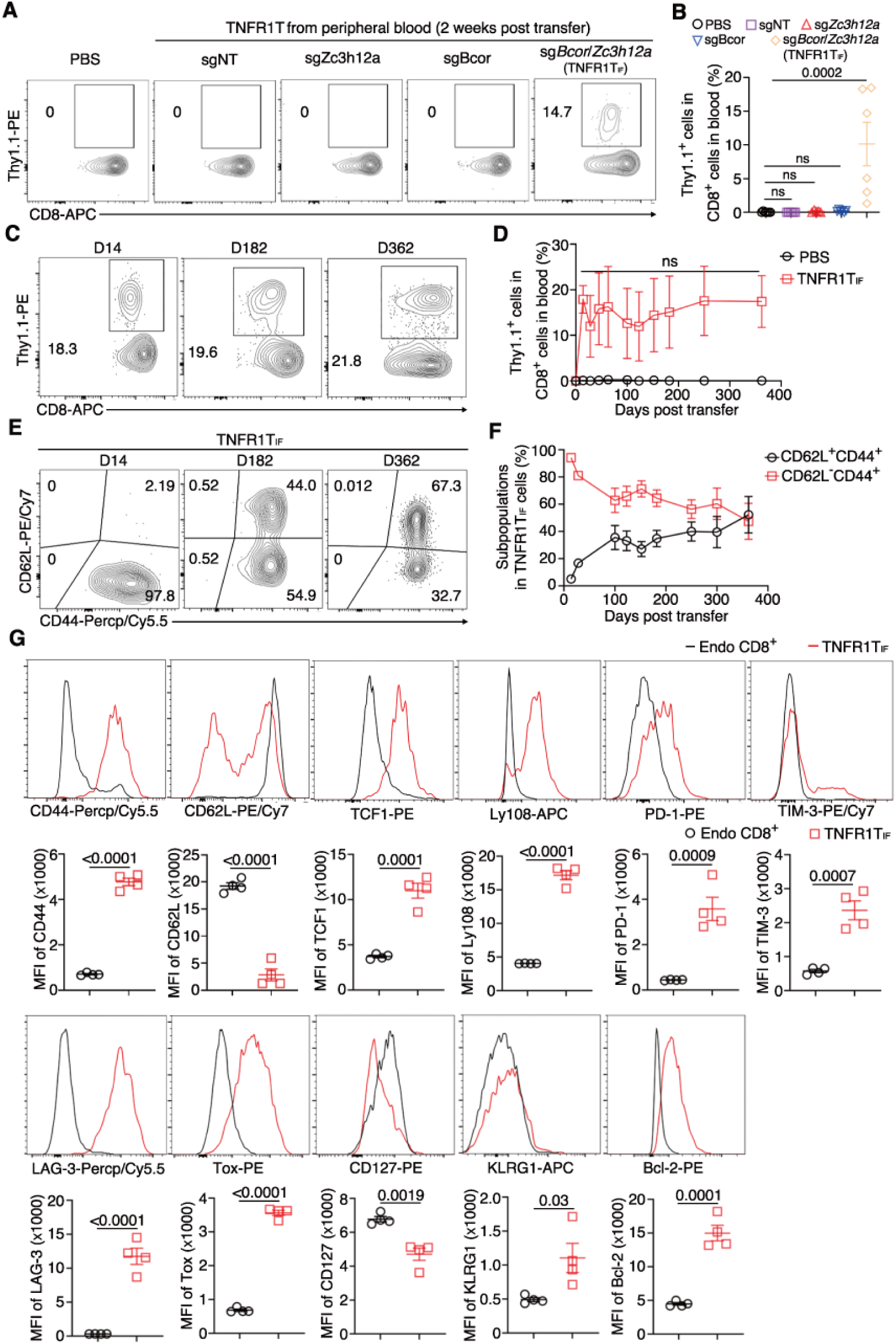
TNFR1T cells lacking BCOR and ZC3H12A (TNFR1T_IF_) expand and persist in immunocompetent mice. **A**, **B**, Percentages of Thy1.1^+^ CAR T cells in peripheral blood two weeks post-transfer. Representative plots (**A**) and statistics (**B**) are shown (n = 6 mice per group). **C**, **D**, Percentages of TNFR1T_IF_ cells in peripheral blood at indicated time points post-infusion. Representative plots (**C**) and statistics (**D**) are shown (n = 4 mice per group). **E**, **F**, Percentages of CD44^+^CD62L^+^ and CD44^+^CD62L^-^ cells within the TNFR1T_IF_ population in peripheral blood at indicated time points post-infusion. Representative plots (**E**) and statistics (**F**) are shown (n = 4 mice per group). **G**, Expression of indicated proteins in TNFR1T_IF_ cells and endogenous CD8^+^ T cells two months post-infusion. Representative plots and statistics are shown (n = 4 mice per group). **B**, **D**, **F, G** Data represent mean ± SEM from one of two independent experiments. *P* values are shown, ns, not significant, one-way ANOVA multiple-comparisons test in (**B**), two-way ANOVA multiple-comparisons test in (**D, F**), two-tailed unpaired Student’s t-test in (**G**).

Given that TNFR1T cells expressing sgNT, sg*Bcor*, or sg*Zc3h12a* exhibited no expansion in vivo (Figure 3A and 3B), our subsequent studies focused on TNFR1T_IF_ cells, with PBS or wild-type TNFR1T cells serving as control (Figure 3A and 3B). Long-term monitoring revealed that TNFR1T_IF_ cells persisted in vivo for at least a year (the end point of the experiment) with stable percentages in peripheral blood (Figure 3C and 3D). Importantly, TNFR1T_IF_ cells could not expand in *Tnf^-/-^*mice devoid of TNF (Figure S1G-S1I), indicating that TNFR1T_IF_ cells respond specifically to TNF in mice.

Two weeks post-infusion, TNFR1T_IF_ cells exhibited a typical effector T cell phenotype (CD44^+^CD62L^-^) (Figure 3E and 3F). Gradually, the percentages of central memory phenotype (CD44^+^CD62L^+^) TNFR1T_IF_ cells increased while effector memory phenotype (CD44^+^CD62L^-^) TNFR1T_IF_ cells diminished (Figure 3E and 3F), suggesting a trajectory resembling memory T cell differentiation. Two months post-infusion, TNFR1T_IF_ cells expressed high levels of CD44, Ly108, and TCF1, along with positivity for PD-1, LAG-3, TOX, as well as low levels of CD127 (Figure 3G). Additionally, a subset of TNFR1T_IF_ cells expressed CD62L and TIM-3 (Figure 3G), suggesting heterogeneity within the TNFR1T_IF_ population at this time point, supporting the notion of continued differentiation of these cells in vivo (Figure 3E and 3F). These characteristics partially align with features of other CAR T_IF_ cells(Jin et al., 2024; Wang et al., 2024), implying that the deficiency of BCOR and ZC3H12A induces a similar program across various CAR T cells.

Together, these data indicate that TNFR1T_IF_ cells are able to expand and persist in fully immunocompetent mice without any conditioning regimen, and their existence in vivo requires the presence of TNF.

### Long-term safety of TNFR1T_IF_ cells

A cohort of mice, infused with either TNFR1T_IF_ cells or PBS, was monitored for up to a year, marking the endpoint of our safety study. Serial sampling of peripheral blood revealed that TNFR1T_IF_ cells persisted in the blood at relatively stable percentages, as illustrated above (Figure 3C and 3D). The presence of TNFR1T_IF_ cells did not affect the percentages of endogenous CD4^+^ and CD8^+^ T cells in peripheral blood (Figure S2A and S2B), nor the activation status of endogenous T cells (Figure S2C and S2D). At 12 months post-infusion, no discernible difference in body weight was observed between mice infused with TNFR1T_IF_ cells and PBS (Figure S2E). Hematoxylin and eosin (H&E) staining of slices from the heart, liver, lung, and kidney did not reveal noticeable differences between mice infused with TNFR1T_IF_ and PBS (Figure S2F). These long-term safety data indicate that the presence of TNFR1T_IF_ cells in mice for up to a year does not obviously affect the health of mice.

### TNFR1T_IF_ cells degrade human TNF in vivo

To investigate whether the depletion of BCOR and ZC3H12A affects the endocytosis of TNF by TNFR1T cells, we transferred TNFR1T_IF_ cells into hTNF-tg mice. TNFR1T_IF_ cells, but not TNFR1T cells, significantly reduced serum hTNF in hTNF-tg mice to near wild-type levels (Figure 4A). Two months post-infusion, TNFR1T_IF_ cells were isolated from mice to perform TNF endocytosis assay described earlier (Figure 1I). Compared with cells kept at 4°C, a significant reduction in RFP-TNF signals was observed in TNFR1T_IF_ cells after incubation at 37°C (Figure 4B and 4C), suggesting degradation or shedding of RFP-TNF. Microscopic imaging of RFP signal showed that RFP-TNF was endocytosed in TNFR1T_IF_ cells after incubation at 37°C (Figure 4D and 4E). These data demonstrate that the absence of BCOR and ZC3H12A in TNFR1T_IF_ cells does not affect their endocytosis of TNF, and TNFR1T_IF_ cells are able to reduce TNF levels in hTNF- tg mice.

**Figure 4.**
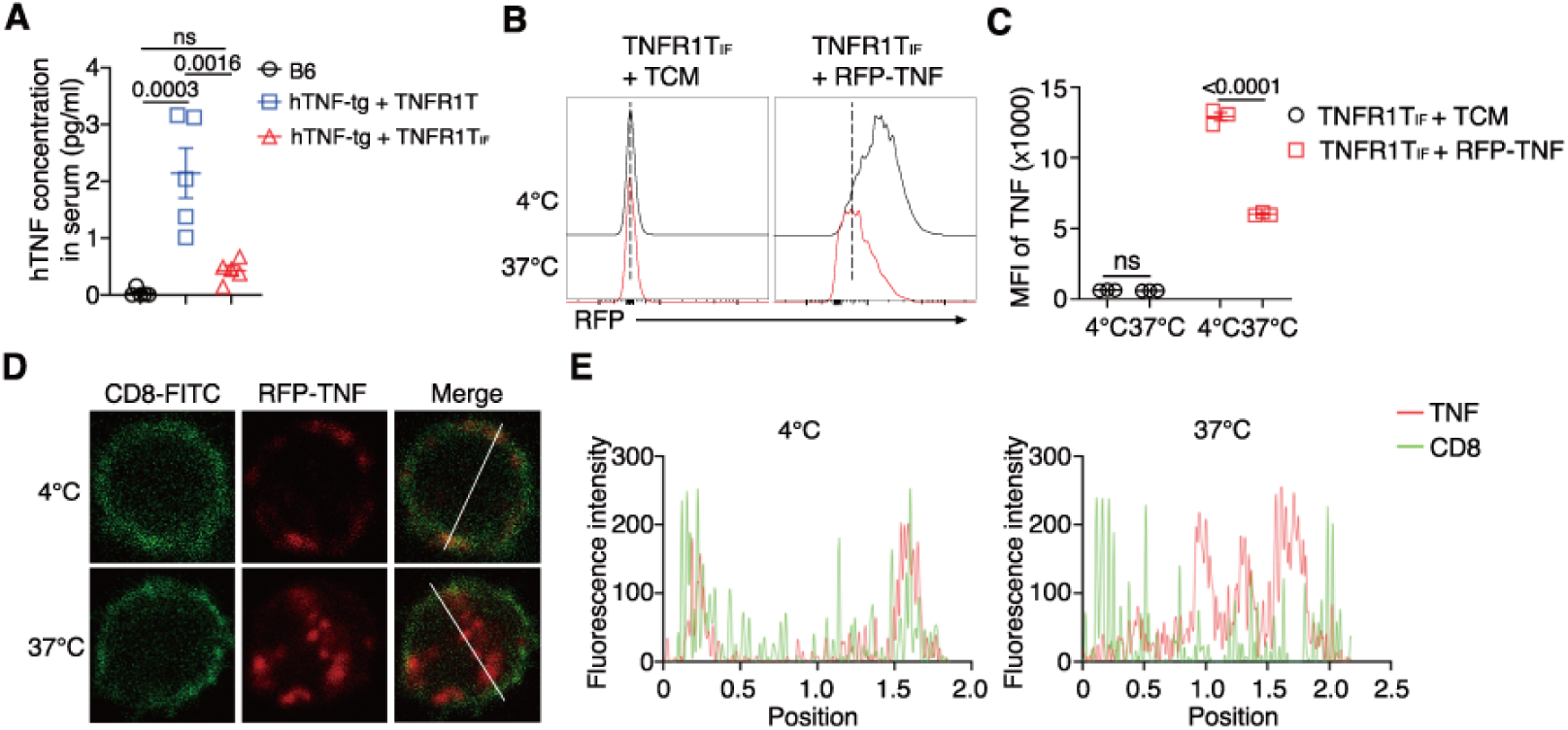
TNFR1T_IF_ cells degrade TNF in vivo. **A**, Serum levels of hTNF in B6 and hTNF-tg mice across treatment groups at 22 weeks of age (n = 5 mice per group). **B**, **C**, TNFR1T_IF_ cells freshly isolated from mice two months post-infusion were analyzed for TNF endocytosis. Flow cytometry analysis of RFP-TNF signals in TNFR1T_IF_ cells was conducted after endocytosis at 37°C for 24 or 48 hours. Representative plots (**B**) and statistics (**C**) are shown (n = 3 independent experiments). **D**, **E**, Immunofluorescent analysis of RFP-TNF localization in TNFR1T_IF_ cells kept at 4°C or after incubation at 37°C for 24 hours. Representative images (**D**) and fluorescence signal quantification along lines drawn in cells (**E**) are shown. **C**, Data represent mean ± SEM from one of three independent experiments. *P* values are shown, ns, not significant, one-way ANOVA multiple-comparisons test.

### TNFR1T_IF_ cells confer long-term remission of RA

Since TNFR1T_IF_ cells degrade hTNF in hTNF-tg mice in vivo (Figure 4), we assessed the impact of TNFR1T_IF_ cells on RA. In female hTNF-tg mice, infusion of TNFR1T_IF_ cells, but not TNFR1T cells, before RA onset prevented RA clinical score (Figure 5A and 5B). TNFR1T_IF_ cells also increased the body weight and grip strength of hTNF-tg mice to near wild-type levels (Figure 5C and 5D). Swelling and deformation of joints in hTNF-tg mice were largely prevented by TNFR1T_IF_ cells, but not TNFR1T cells (Figure 5E and 5F). In male hTNF-tg mice, infusion of TNFR1T_IF_ cells before RA onset also prevented RA clinical score and increased grip strength (Figure S3A- S3C). These data demonstrate that TNFR1T_IF_ cells prevent RA development if infused before disease onset.

**Figure 5.**
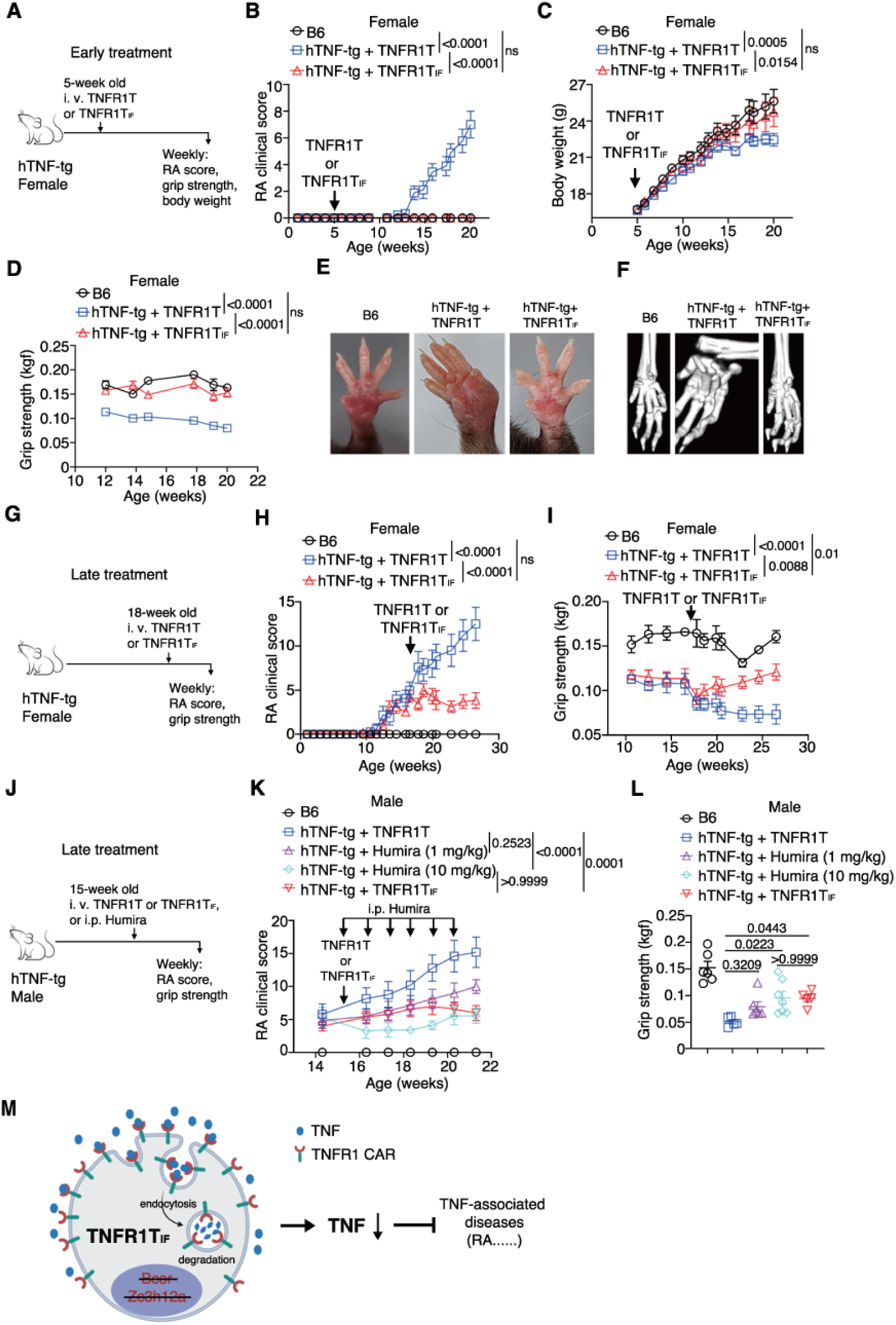
A single infusion of TNFR1T_IF_ cells confers long-term remission of RA in hTNF-tg mice. **A**, Experimental design for early RA treatment in female hTNF-tg mice. **B**, RA clinical scores in hTNF-tg and B6 mice across treatment groups (n = 9 mice per group). **C**, Grip strength in hTNF-tg and B6 mice across treatment groups (n = 9 mice per group). **D**, Body weights of hTNF-tg and B6 mice across treatment groups (n = 9 mice per group). **E**, Representative images of front paws from hTNF-tg and B6 mice at 20 weeks of age across treatment groups. **F**, Representative μCT images of front paws from hTNF-tg and B6 mice at 20 weeks of age across treatment groups. **G**, Experimental design for late RA treatment in female hTNF-tg mice. **H**, RA clinical scores in hTNF-tg and B6 mice across treatment groups (n = 5 mice in B6 group, n = 10 mice in hTNF-tg + TNFR1T group, n = 9 mice in hTNF-tg + TNFR1T_IF_ group). **I**, Grip strength in hTNF-tg and B6 mice across treatment groups (n = 5 mice in B6 group, n = 10 mice in hTNF- tg + TNFR1T group, n = 9 mice in hTNF-tg + TNFR1T_IF_ group). **J**, Experimental design for late RA treatment in male hTNF-tg mice. **K**, RA clinical scores in hTNF-tg and B6 mice across treatment groups (n = 6 mice in B6 group, n = 5 mice in hTNF-tg + TNFR1T group, n = 6 mice in hTNF-tg + 1 mg/kg Humira group, n = 7 mice in hTNF-tg + 10 mg/kg Humira group, n = 5 mice in hTNF-tg + TNFR1T_IF_ group). **L**, Grip strength in hTNF-tg and B6 mice across treatment groups (n = 6 mice in B6 group, n = 5 mice in hTNF-tg + TNFR1T group, n = 6 mice in hTNF-tg + 1 mg/kg Humira group, n = 7 mice in hTNF-tg + 10 mg/kg Humira group, n = 5 mice in hTNF-tg + TNFR1T_IF_ group). **M**, Proposed model of TNFR1T_IF_ cells degrading TNF to treat TNF-associated diseases. **B**, **C**, **D**, **H**, **I**, **K**, **L**, Data represent mean ± SEM from one of two independent experiments. *P* values are shown, ns, not significant, two-way ANOVA multiple-comparisons test in in (**B**, **C**, **D**, **H**, **I**, **K**), one-way ANOVA multiple-comparisons test in (**L**).

We next infused TNFR1T_IF_ cells into male hTNF-tg mice after the onset of RA to assess their therapeutic efficacy (Figure S3D). Under this protocol, TNFR1T_IF_ cells gradually reduced the RA clinical score and partially improved grip strength (Figure S3E and S3F), indicating efficacy even after RA onset in male hTNF-tg mice. Furthermore, when TNFR1T_IF_ cells were administered to 18-week-old hTNF-tg mice with fully developed RA, they still suppressed RA clinical score and enhanced grip strength (Figure 5G-5I).

Finally, we performed a side-by-side comparison of TNFR1T_IF_ cells and Humira in hTNF-tg mice with RA (Figure 5J). While a low dose of Humira (1 mg/kg, weekly) showed limited efficacy, a single infusion of TNFR1T_IF_ cells achieved comparable efficacy to a high dose of Humira (10 mg/kg, weekly) in RA management (Figure 5K and 5L). Collectively, these data demonstrate that a single infusion of TNFR1T_IF_ cells provides durable therapeutic efficacy for RA in hTNF-tg mice, similar to repeated high-dose Humira treatment.

## Discussion

While traditional CAR T cells eliminate cells, the CAR T cells described in this study “eliminate” proteins. TNFR1T_IF_ cells represent the first CAR T cells targeting a soluble factor for degradation, marking significant conceptual and technological advancements. All known TPD technologies depend on the engagement of host protein degradation machinery, which cause unforeseeable side effects(Duyvendak et al., 2005; Hamadani et al., 2007; Merrill et al., 2022; Richardson et al., 2023; Sohlbach et al., 2006). In contrast, TNFR1T_IF_ cells degrade TNF via exogenously introduced CAR T cells, bypassing the host protein degradation machinery and reducing the likelihood of affecting host physiology.

All traditional drugs must be administered repeatedly to maintain efficacy. In contrast, TNFR1T_IF_ cells, as a living drug, exhibits unprecedented durability, with a single infusion conferring long-term degradation of TNF and sustained remission of RA. This offers a potential functional “cure” for a prevalent chronic inflammatory disease that typically requires repeated drug administration over years or even decades, heralding a paradigm shift in chronic disease management.

Mechanistically, TNFR1T_IF_ cells achieves TNF degradation through TNFR1 CAR-mediated endocytosis of TNF (Figure 5M). Since CARs can undergo partial recycling back to the cell surface or be synthesized de novo by CAR T cells(Li et al., 2020), TNFR1T_IF_ cells are able to continuously degrade TNF. This explains why TNFR1T_IF_ cells potently diminished serum hTNF to near wild-type levels in hTNF-tg mice, which constitutively produce hTNF from a transgene, a condition much more severe than that of most RA patients in clinical settings. Additionally, TNFR1T_IF_ cells circumvents side effects associated with protein drugs, such as injection site reactions, anaphylaxis, and the generation of anti-drug antibodies.

To overcome the challenge of TNFR1T cells’ inability to expand in vivo, we implemented a recently reported double knockout of BCOR and ZC3H12A to enhance expansion and persistence in mice(Wang et al., 2024), resulting in TNFR1T_IF_ cells. The mechanism underlying the long-term persistence of TNFR1T_IF_ cells in vivo remains incompletely investigated in this study due to the complete absence of all control CAR T cells, including wild-type, BCOR-deficient, and ZC3H12A-deficient TNFR1T cells (Figure 3A and 3B). These control cells are essential for comprehensive mechanistic assays such as ATAC-seq, RNA-seq, and others. Nevertheless, TNFR1T_IF_ cells share certain features with other BCOR- and ZC3H12A-deficient CAR T cells, suggesting that BCOR and ZC3H12A deficiencies induce a similar program across different CAR T cells. Future studies will delve into the mechanisms underlying the induction and maintenance of TNFR1T_IF_ cells. Despite these limitations, our data conclusively demonstrate the safety, efficacy, and mechanism of TNF degradation, laying a foundation for clinical studies.

In theory, alternative methods capable of promoting CAR T cell expansion and persistence in immunocompetent mice without the need for conditioning regimen might be applicable to implant TNFR1T cells into mice. However, as of the manuscript submission, we are not aware of such alternative methods. A significant obstacle in extending CAR T cell therapy to non-cancerous diseases lies in the requirement for a pre-treatment conditioning regimen, often involving intense chemotherapeutic measures such as cyclophosphamide and/or fludarabine(Muranski et al., 2006; Restifo et al., 2012). In the context of non-cancerous diseases, the severe side effects associated with chemotherapeutic conditioning such as cytokine release syndrome, heightened infection risks due to immunodeficiency, and genome instability resulting from genotoxicity, may not justify the application of CAR T cells.

Our study marks a pivotal shift in the targets of CAR T cells, transitioning from cells to soluble factors. Beyond TNF, numerous other inflammatory cytokines implicated in many diseases, such as IL-1β, IL-4, and IL-17, stand as potential candidates for targeting by CAR T cells using the strategies outlined here. Expanding beyond soluble factors, extracellular protein aggregates, such as Aβ aggregates in Alzheimer’s disease(Karran and De Strooper, 2022), could also become potential targets for CAR T cells.

In summary, our study introduces a novel platform for redirecting CAR T cells to target extracellular soluble factors for degradation. This advancement not only broadens the spectrum of treatable diseases for CAR T cells but also provides a lasting therapeutic option for chronic diseases that require repeated drug administration.

## Materials and mèthods Mice and cell lines

TNF knockout (*Tnf^-/-^*) mice on C57BL/6 background were provided by Dr. Xin Lin (Tsinghua University). C57BL/6 (The Jackson Laboratory, Cat# JAX:000664, RRID:IMSR_JAX:000664), hTNF transgenic mice (C57BL/6 background) (Taconic Biosciences, Cat#1006), *Tnf^-/-^*mice and Cas9 transgenic mice (The Jackson Laboratory, Cat# JAX:026430, RRID:IMSR_JAX:026430) were housed in a specific pathogen-free environment at Tsinghua University’s Laboratory Animal Research Center in Beijing, China. The facilities for these animals were approved by the Beijing Administration Office of Laboratory Animal. All animal-related procedures were sanctioned by the Institutional Animal Care and Use Committee (IACUC).

Phoenix-Eco cells (Cell Biolabs, Cat# RV-101) and HEK293T cells (ATCC, Cat# CRL-3216, RRID:CVCL_0063) were grown in Dulbecco’s Modified Eagle Medium (DMEM, Gibco), supplemented with 5% fetal bovine serum (FBS), 2 mM glutamine, and antibiotics (penicillin and streptomycin). Cells were maintained in a humidified incubator at 37 °C. All cell lines were tested for mycoplasma contamination using the TransDectTM PCR Mycoplasma detection Kit (TRAN, FM311) and were confirmed to be negative for mycoplasma.

### Plasmids

The TNFR1 CAR consisted of mouse TNFR1 ectodomain, mouse CD28 transmembrane and signaling domain, followed by mouse CD3ζ. This TNFR1 CAR was inserted into the pMSCV- hU6-sgRNAs-EFS-Thy1.1-CAR backbone. For dual sgRNA expression, another U6 promoter was inserted. The sequences of sgRNAs: sgNon-targeting (sgControl), 5′- TTCGCACGATTGCACCTTGG-3′; sg*Zc3h12a*, 5′-CTAGGGGAATTGGTGAAGCA-3′; sg*Bcor*, 5′-ACTGG GCAATACCGCAACAG-3′.

### Antibodies

Anti-mouse Thy1.1 (OX-7), biotin, Cat# 202510, BioLegend, RRID:AB_2201417; anti-mouse Thy1.1 (OX-7), PE, Cat# 202524, BioLegend, RRID:AB_1595524; anti-mouse TNFR1 (55R- 286), PE, Cat# 113003, BioLegend, RRID:AB_313532; anti-mouse CD8α (53-6.7), APC, Cat# 100712, BioLegend, RRID:AB_312751; anti-mouse CD8α (YTS156.7.7), FITC, Cat# 126606, BioLegend, RRID:AB_961295; anti-mouse CD4 (RM4-5), APC/Cyanine7, Cat# 100526, BioLegend, RRID:AB_312727; anti-mouse CD69 (H1.2F3), PE, Cat# 104508, BioLegend, RRID:AB_313111; anti-mouse CD44 (1M7), PerCP/Cyanine5.5, Cat# 103032, BioLegend, RRID:AB_2076204; anti-mouse CD62L (MEL-14), PE/Cyanine7, Cat# 104418, BioLegend, RRID:AB_313103; anti-mouse TCF1 (C63D9), PE, Cat# 14456, Cell Signaling Technology, RRID:AB_2798483; anti-mouse PD-1 (29F.1A12), PE, Cat# 135206, BioLegend, RRID:AB_1877231; anti-mouse TIM-3 (RMT3-23), PE/Cyanine7, Cat# 25-5870-82, Thermo Fisher Scientific, RRID:AB_2573483; anti-mouse LAG-3 (C9B7W), PerCP-eFluor™ 710, Cat# 46-2231-82, Thermo Fisher Scientific, RRID:AB_11151334; anti-mouse Bcl-2 (BCL/10C4), PE, Cat# 633508, BioLegend, RRID:AB_2290367; anti-mouse KLRG-1 (2F1), APC, Cat# 561620, BD Biosciences, RRID:AB_10895798; anti-mouse CD127 (A7R34), PE, Cat# 135010, BioLegend, RRID:AB_1937251; anti-mouse Ly108 (13G3-19D), APC, Cat# 17-1508-82, Thermo Fisher Scientific, RRID:AB_10717668; anti-mouse Tox (TXRX10), PE, Cat# 12-6502-82, Thermo Fisher Scientific, RRID:AB_10855034; Streptavidin, PE, Cat# 405204, BioLegend; Streptavidin, APC, Cat# 405243, BioLegend; Streptavidin, APC/Cyanine7, Cat# 405208, BioLegend; anti-mouse B220 (RA3-6B2), eFluor 450, Cat# 48-0452-82, BioLegend, RRID: AB_1548761; anti-mouse CD11b (M/70), PerCP/Cy5.5, Cat# 45-0112-82, Thermo Fisher Scientific, RRID:AB_953558; anti-mouse TCRβ (H57-597), APC, Cat# 17-5961-82, Thermo Fisher Scientific, RRID:AB_469481.

### Retrovirus production

Retroviruses were packaged by co-transfection of Phoenix-Eco cells with indicated plasmid and helper plasmid pCL-Eco (Addgene #12371) using calcium phosphate precipitate-mediated transfection. The viral supernatant was collected at 48 hours and 72 hours post-transfection, filtered through 0.45 mm filters, aliquoted, and frozen at -80 °C.

### Mouse T cells culture, viral transduction and adoptive transfer

CD8^+^ T cells were isolated by a negative selection Kit (BioLegend, Cat# 480035) from spleens and lymph nodes of Cas9^+^ mice, then activated overnight with 1 μg/ml anti-CD3 (BioXCell, BE0001-1, RRID:AB 1107634) and 1 μg/ml anti-CD28 (Bio X Cell, BE0015-1, RRID:AB_1107624) in T cell medium (TCM): RPMI1640 medium (Gibco) supplemented with 5% fetal bovine serum, 2 mM glutamine, 55 mM b-mercaptoethanol, 1 mM sodium pyruvate, 100 units/ml penicillin, 100 mg/ml streptomycin and 2 ng/ml IL-2 (PeproTech, Cat# 200-02-1000).

The following day, activated T cells were spin-infected at 2,000 g with retrovirus described above at 33 °C for 2 hours. Then, cells were washed and cultured in fresh TCM with IL-2. Twenty-four hours after spin-infection, the efficiency of transduction was determined by flow cytometry. All adoptive transfers to indicated mice (including C57BL/6 mice, *Tnf^-/-^* mice, and hTNF transgenic mice) were performed at 24 hours post-transduction via tail vein in the absence of any conditioning regimen.

### RA model

The hTNF-tg mice spontaneously developed severe RA, similar to human RA. These mice exhibited arthritis in both fore and hind paws starting around 8-12 weeks of age, with the condition worsening as they aged. TNFR1T cells, TNFR1T_IF_ cells or Humira were transferred into hTNF-tg mice via tail vein at indicated time points. Clinical signs of hTNF-tg mice including RA clinical scores, body weight, and grip strength were monitored weekly. The scoring criteria for toes were as follows: 0, no signs; 1, mild deformity; 2, severe deformity or swelling. The scoring criteria for wrists and ankles are as follows: 0, no signs; 1, mild swelling; 2, severe swelling; 3, severe swelling along with noticeable distortion; 4, severe swelling along with severe distortion. The scores of all 4 paws of a mouse were combined and used as the total RA score.

The grip strength of both the forelimbs and hindlimbs was evaluated using a grip strength meter. As the mouse grasped the bar, the peak pull force was recorded by a digital force transducer. The highest score from three consecutive trials was used to determine the grip strength of the forelimbs and hindlimbs.

### Measurement of hTNF

To measure hTNF levels in serum, a cytometric bead array (CBA) kit (BD Biosciences, Cat# 551811) was used. Blood samples were collected from mice in 1.5 ml sterile tubes, left at room temperature for 20 minutes, and then centrifuged at 2,000 rpm at 4°C for 10 minutes. The supernatants were carefully transferred to new tubes and centrifuged again at 2,000 rpm at 4°C for another 10 minutes. The final supernatants (serum) were promptly subjected to CBA assay to the protocol provided by the Kit.

### Micro-computed tomography (μCT)

Bone samples were preserved in a solution of 4% paraformaldehyde (PFA). μCT images of the PFA-fixed bone samples were captured using a Quantum GX μCT (PerkinElmer, Inc., USA) at 60 kV and 160 μA. Images were taken with a 9 μm voxel size resolution and a 72 x 72 mm field of view (FOV) in high-resolution mode. The captured images were visualized and reconstructed using the built-in software within the Quantum GX system.

### Histology

Tissues from euthanized animals were fixed in 4% PFA and embedded in paraffin. 5-µm sections were stained with hematoxylin and eosin (H&E).

### Flow cytometry

Single-cell suspension was prepared from indicated tissues/organs. Surface proteins were stained with antibodies in the presence of Fc block in FACS buffer (PBS containing 1% FBS, 2 mM EDTA) at 4 °C for 15 min. Intracellular staining of cytoplasmic and nuclear proteins was performed with Transcription Factor Staining Buffer kit (BD Pharmingen). Dead cells were excluded by DAPI staining or LIVE/DEAD Fixable Near-IR Dead Cell Stain Kit (Cat# L34976, Invitrogen). Samples were analyzed by an LSR Fortessa cytometer (BD). Flow cytometry data were analyzed using Flowjo software. Cell sorting was performed on a S3e cell sorter (Bio-Rad).

### TNF internalization assay

TNFR1T cells or TNFR1T_IF_ cells were first labelled with RFP-TNF fusion protein (expressed and purified from HEK293T cells in house) or anti-TNF antibody on ice for 20 minutes. After washing out unbound RFP-TNF and anti-TNF antibody, CAR T cells were resuspended in TCM and split into two parts. One part of cells was kept at 4°C, and the other was put into a 37°C incubator to allow endocytosis for 24 hours. Subsequently, cells were stained with indicated antibodies at 4°C for 15 minutes. The signals of RFP-TNF and anti-TNF antibody were examined by FACS.

To visualize the intracellular localization of RFP-TNF, cells underwent the above treatment were loaded on a glass slide coated with poly-lysine. After fixation, the slides were mounted with antifade mountant (Thermo Fisher). Images of the cells were captured using a Zeiss LSM780 microscope and analyzed using ImageJ software.

### Quantifications and statistical analysis

The statistical information of each experiment, including the statistical methods, the *P* value and sample numbers (n) were shown in figure or figure legends. GraphPad Prism 9 (https://www.graphpad.com) was used to plot all graphs and to perform statistical and quantitative assessments. Error bars represent standard error of mean (SEM).

## Data availability

Source data will be provided upon request. All other data that support the findings of the present study are present in the article or are available from the corresponding author upon request.

## Acknowledgments

This research was supported by National Natural Science Foundation of China (grant 82350108 to M. P.), Vanke Special Fund for Public Health and Health Discipline Development Tsinghua University (NO.2022Z82WKJ013, to M. P.), Tsinghua University DUSHI Program

(52302102323, to M.P.), Tsinghua-Peking Center for Life Sciences (to M. P.), SXMU-Tsinghua Collaborative Innovation Center for Frontier Medicine (to M. P.).

## Author contributions

Y.W. performed experiments and analyzed data; N.Y. provided technical help and supervised the project; M.P. conceived and supervised the project, analyzed and interpreted data, and wrote the manuscript with inputs from all authors.

## Competing interests

A patent application has been filed based on findings described in this study.

---

**Figure S1.**
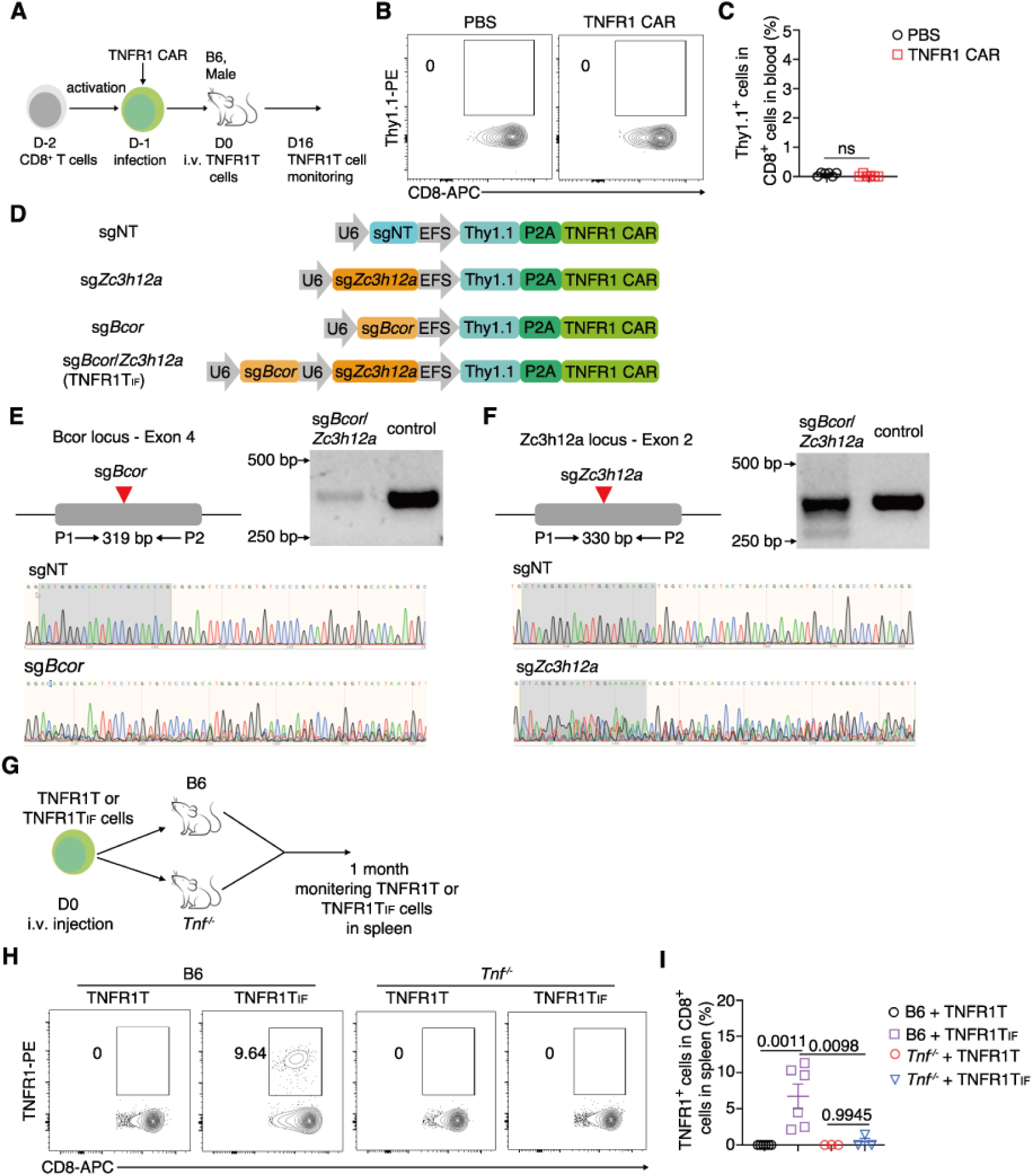
Expansion of TNFR1T and TNFR1T_IF_ cells in B6 and *Tnf^-/-^* mice. **A**, Experimental design for assessing TNFR1T cell expansion in B6 mice. **B**, **C**, Percentages of Thy1.1^+^ CAR T cells in the peripheral blood 16 days post-transfer. Representative plots (**B**) and statistics (**C**) are shown (n = 6 mice per group). **D**, Schematic of a single-vector system for TNFR1 CAR expression along with the indicated sgRNA. **E**, **F**, Editing efficiency of *Bcor* (**E**) and *Zc3h12a* (**F**) was assessed by DNA sequencing. Cleavage sites are marked by red arrows. PCR primers spanning the cleavage site were used to amplify the target genomic region. The PCR products, covering the regions with sgRNA cleavage sites, are displayed along with representative sequencing tracks. **G**, Experimental design for assessing TNFR1T_IF_ cell expansion in B6 and *Tnf^-/-^*mice. **H**, **I**, Percentages of TNFR1^+^ CAR T cells in the spleen one-month post-transfer. Representative plots (**H**) and statistics (**I**) are shown (n = 6 mice in B6 group, n = 3 mice in *Tnf^-/-^*group). **C**, **I**, Data represent mean ± SEM from one of two independent experiments. *P* values are shown, ns, not significant, two-tailed unpaired Student’s t-test in (**C**), one-way ANOVA multiple-comparisons test in (**I**).

**Figure S2.**
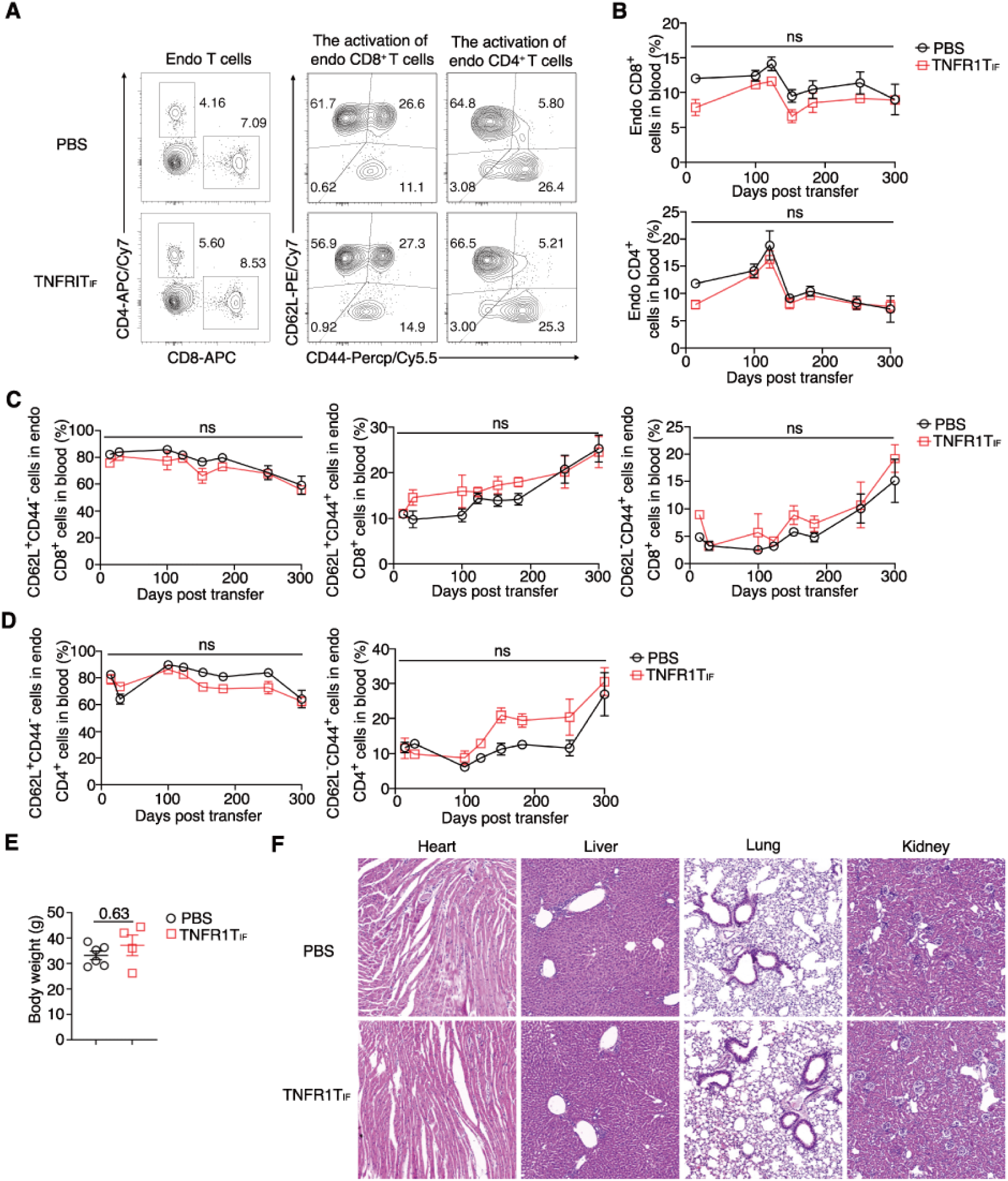
Long-term safety of TNFR1TIF cells in B6 mice. A, B, FACS analysis of endogenous T cells in peripheral blood from mice infused with PBS or TNFR1T_IF_ cells. Representative plots (**A**) and statistics (**B**) are shown (n = 6 mice in PBS group, n = 4 mice in TNFR1T_IF_ group). **C, D,** FACS analysis of activation status in endogenous CD4^+^ T cells (**C**) and CD8^+^ T cells (**D**) from peripheral blood of mice infused with PBS or TNFR1T_IF_ cells. Representative plots (**C**) and statistics (**D**) are shown (n = 6 mice in PBS group, n = 4 mice in TNFR1T_IF_ group). (**E**) Body weight of mice infused with PBS or TNFR1T_IF_ cells at 12 months post-infusion (n = 6 mice in PBS group, n = 4 mice in TNFR1T_IF_ group). (**F**) Hematoxylin and Eosin (H&E) staining of slices of heart, liver, lung, and kidney tissue sections from mice infused with PBS or TNFR1T_IF_ cells, 12 months post-infusion. Representative images are shown. *P* values are shown, ns, not significant, two-way ANOVA multiple-comparisons test.

**Figure S3.**
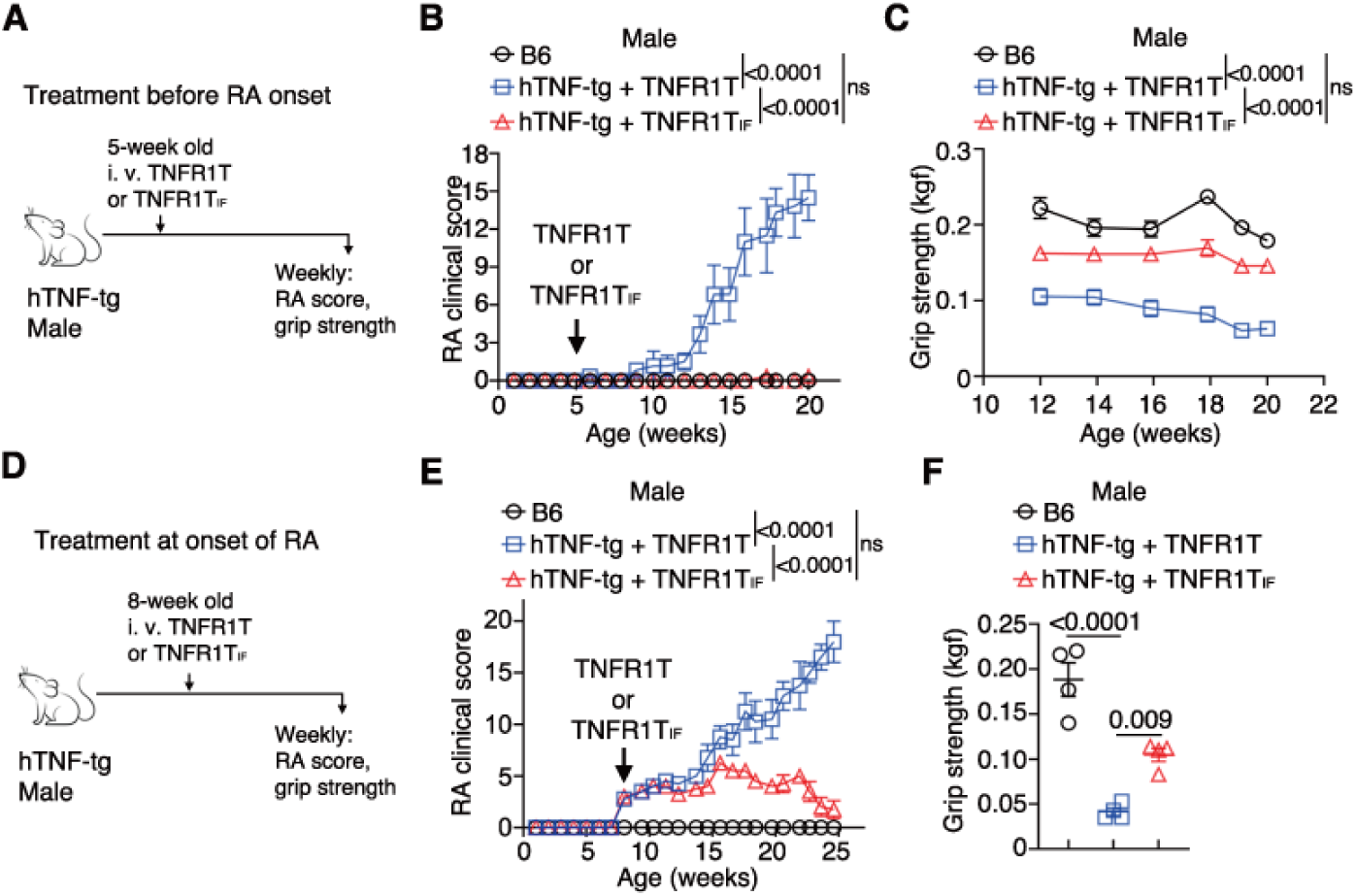
Efficacy of TNFR1T_IF_ cells for RA in hTNF-tg mice. **A**, Experimental design for early RA treatment in male hTNF-tg mice. **B**, RA clinical scores in hTNF-tg and B6 mice across treatment groups (n = 6 mice per group). **C**, Grip strength in hTNF-tg and B6 mice across treatment groups (n = 6 mice per group). **D**, Experimental design for RA treatment starting from disease onset in female hTNF-tg mice. **E**, RA clinical scores in hTNF-tg and B6 mice across treatment groups (n = 4 mice per group). **F**, Grip strength in hTNF-tg and B6 mice at 23 weeks of age across treatment groups (n = 4 mice per group). **B**, **C**, **E**, **F**, Data represent mean ± SEM from one of two independent experiments. *P* values are shown, ns, not significant, two-way ANOVA multiple-comparisons test in (**B**, **C**, **E**), one-way ANOVA multiple-comparisons test in (**F**).

